# A new idea on the evolution of biodiversity

**DOI:** 10.1101/019828

**Authors:** Roberto Cazzolla Gatti

## Abstract

The understanding of the mechanisms that allow the origin of species has shed light on many processes such as speciation, adaptation and extinction, but how to explain the existence of such a huge diversity of species on Earth still remains a mystery. Many theories and evidence have corroborated the processes that allow species to evolve or become extinct. Currently there are different hypotheses but no clear demonstrations of the factors that maintain the species diversity of ecosystems. Those based on competitive principles have been criticized from both theoretical and empirical approaches. Only by studying biodiversity in the context of evolution, natural history and ecology (taking into consideration the *avoidance of competition* and dispersal abilities, the phenotypic plasticity, the heterogeneous landscape, the facilitation and the *endogenosymbiosis*) we can understand values and gaps of past theories trying to provide a broader understanding of life on Earth towards a Unified Theory of Biodiversity.

## Introduction

One of the greatest scientific revolutions in human history is the theory of evolution proposed by Charles Darwin (^1^) and, at the same time, by Alfred R. Wallace (^2^). The understanding of the mechanisms that allow the origin of species and explain their current presence on planet Earth has shed light on many processes such as speciation, adaptation and extinction. In the following decades many hypotheses and evidence have corroborated the mechanisms allow species to evolve, coexist, compete, cooperate or become extinct. Certainly one of the most debated aspect of the hypotheses which have been developed (^3^,^4^) is actually finding out which factors allow species to coexist in a given time within the same environment. Starting from the principle of competitive exclusion of Gause (^5^, ^6^) until Connell’s ghost of competition in the past (^7^), the importance of intra-and inter-specific competition for the evolution of biodiversity has been stressed. These theories and hypotheses suggest that competition tends to differentiate the ecological requirements after repeated interactions and allow the presence of many different species in the same area. Recently, the principles based on competitive reasons for the explanation of biodiversity have been criticized from both theoretical and empirical approaches. Only by studying biodiversity in the context of evolution and ecology we can understand the values and the flaws of past theories and provide a broader understanding of life on Earth. The path towards a Unified Theory of Biodiversity starts from this consideration.

The concept of the niche as the set of ecological requirements, from the reproductive to the alimentary ones, developed by Charles Elton (^8^) and improved by Evelyn Hutchinson (^9^) with the definition of hyper-volume, is a powerful tool for understanding the role of each species in its environment. To date, however, a niche has been often considered as something static and unchanging (^10^), while the dynamics and feedback loops that allow their creation and the interaction between those of different species have not been carefully analyzed. Furthermore, most studies have focused on the role of species in the environment and have frequently neglected the understanding of the role that environment and space have on the dynamics of species and the formation of niches (^11^).

Darwin expressed is idea on the origin of species using the following statements: severe competition between conspecifics and close varieties drives evolution and diverging evolution leads to less intense competition and coexistence of many species. The weakening competition with ecological differentiation was an unavoidable element of Darwin’s theory. He needed it to interpret evolution through selection and the huge species diversity he observed within a common conceptual framework (^12^). Unfortunately this paradigm has now been criticized by several theories and empirical evidence (^26,27,28,29,30^). For Darwin, the unavoidable limit of population growth and intraspecific competition were both essential for the struggle for existence.

Interspecific competition or competition between different populations becomes weaker when they are checked by different factors or resources. Considering Gause’s principle and the competitive Lotka-Volterra (^13^,^14^) in an ecological framework these models express the essence of the Darwinian paradigm.

On the other hand, starting from 1961 Hutchinson’s provocation of the “paradox of plankton” (^15^) a series of topics using different approaches has been proposed to explain why the principle of competitive exclusion is not found in “nature”. The reason probably lies in the fact that ecologists have not questioned some of the principles of natural selection. Most ecological models are too simplistic and are often considered as outdated. Here I propose a new model that tries to collect the new trends in biodiversity and evolutionary science, pointing out the importance of “avoidance of competition, biological history, “*endogenosymbiosis*” and “ three-dimensionality as the main forces that structure ecosystems.

## The effect of intraspecific competition

Charles Darwin’s Origin of Species suggested that the struggle for existence was the main driver of the evolution of species and indicated the survival of the strongest as an evidence of adaptation to the environment by individuals. Intraspecific competition is a mechanism that has been considered for a long time as fundamental to explain the coexistence of species, which reduces the resources available to individuals of their own species allowing others to survive in the same environment. This competition can occur for exploitation of food resources, space or indirectly interference and reducing the availability of food, space or the possibility of meeting conspecifics, it guarantees a certain level of diversity. These interactions are generally referred as density-dependent and the oscillation in the levels of birth rate and mortality within populations as the consequence. Achieving a balance between birth rates and mortality, in accordance with the external environment, has been defined by the term *carrying capacity*, below which the birth rate exceeds mortality and the population grows, while above it the opposite occurs.

The carrying capacity, however, should not be seen as a static threshold, but rather as a dynamic range of density that regulates the number of individuals in populations. Without the constraint of this ecological edge the population would increase following an exponential growth, with an intrinsic rate of natural growth. In real conditions, however, sooner or later models of population growth assume a sigmoid logistical shape, reaching levels of carrying capacity.

We can identify two main factors which, while helping to minimize the effects of intraspecific competition, also ensure the coexistence of species. The first concerns the spatial distribution. It is well known that essentially all species tend to have dispersion and migration strategies that enable individuals belonging to the same population to limit competition for resources and space. MacArthur and Wilson (^16^) in their equilibrium theory of island biogeography stressed the importance of immigration and emigration in the total amount of the number of species of a given territory.

The second factor proposed to explain the coexistence of species, which I will discuss in depth below, being the main focus of my discussion, is biological history. In an evolutionary context, each species derives from the speciation of its ancestor and its role, functions and niche in the present derive from the interactions of its phylogenetic line in the past. The importance of the trade-off between growth and reproduction, competition and cooperation, speciation and extinction plays a major role in understanding the patterns of the evolution of biodiversity.

## Interspecific competition and the struggle among species

Interspecific competition is probably the interaction between species that has been most studied by ecologists. Several mathematical models have been proposed to define the relationships between different species that inhabit a particular environment. Most research have focused either on small ecosystems or on those reproduced in laboratories that are easily controllable. Some have analyzed the interactions between different species of diatoms (^17^), grassland plants (^18^), barnacles (^19^), bumblebees (^20^), rodents (^21^), etc. From these studies some general observations have been suggested: 1) the spatial scale is crucial for the analysis of coexistence because some species seem to coexist on a large spatial scale but, at a lower resolution, they display distinct distributions; 2) experimental ecosystems tend to be too simplified compared to natural ones and therefore they may lead to erroneous results; 3) some species that might survive in a given territory without competition may be excluded by interspecific competition; and 4) the “fundamental niche” of a species is the ensemble of all their vital functions that allow them to live in some environments where other species are not able to. According to these results in nature the niche becomes “realized” only because other species, that may trigger competition, are there as well. Thus it seems that species can coexist when each of them has a fundamental and, so realized, niche space in its environment. Here I show that there are other ways in which fundamental niches can be developed and filled.

All the above observations led to the synthesis in the principle of competitive exclusion, which shows that the coexistence of two species in a stable environment is likely only if they have occupied different niches. Should they completely overlap, one of two competing species will eliminate the other. This principle was mathematically modelled by Vito Volterra and James Lotka. In spite of the validity of the model and formal proof of Gause’s law, the Lotka-Volterra system shows several flaws in analyzing the dynamics of real ecosystems. First, it uses a coefficient of competition that takes into account the interaction between just two species. This rarely occurs in nature where we rather find networks of relationships between species. Even if there are several multispecies extensions of Lotka-Volterra system (^22^;^23^) they all use the same simplified equations that does not actually match with reality. Moreover, it does not prove that the species in nature are really competing with one another or that they ever did it in the past. A closer look at all the species living in a particular environment will show that each of those has a unique realized niche. No scientific evidence of the real competition among species has been shown yet. Again, as for the case of intraspecific competition, the effect of the biological history on the dynamics that shape biodiversity seems to be of paramount importance. I will look into this aspect, as in the previous case, in the central part of the discussion, but here it is necessary to focus over one point: although the species cannot compete in the present, it is likely that their ancestors had some type of interaction in the past that led to a differentiation of niches which today allows them to coexist.

Moreover, unlike what has been shown by the principle of competitive exclusion and often created in laboratory experiments, environmental heterogeneity and variability are important aspects of most of the habitats and ensure that the conditions for competition between species do not occur or are insignificant.

Another problem that seems to be still only partially resolved is whether the evolutionary effects, “of the past”, are more important than the ecological ones, “in the present”, in reducing interspecific competition and allowing coexistence. I will show that the theoretical construct of my discussion and the scientific evidence move towards both an evolutionary explanation for the structure of ecosystems and an ecological one for maintaining the relationships that have been settled. Just to make a well-known example, the case of Darwin’s finches of *Geospiza* genus, examined by the English naturalist at Galapagos, shows a differentiation of niches as a result of evolutionary forces that have pushed their beaks to adapt to different types of food following a process known as character displacement, which tends to amplify small phenotypic differences in the long term.

## Is the neutrality of species niches necessary?

Stephen P. Hubbell proposed about ten years ago the unified neutral theory of biodiversity and biogeography UNTB (^24^), which assumes that all species belonging to the same trophic level of an ecological community are “neutral” in relation to their fitness. This implies that there are not real differences between the niches of each species and that their success is dictated by the randomness of the moment. Hubbell assumes that population densities are constant and that the most common species, having a higher birth rate, are more likely to speciate than rare ones. Although this theory predicts very well the distribution of species by means of a dynamic balance of immigration, speciation and extinction, its main merit seems to have reassessed the importance of niches in the maintenance of biodiversity (^25^). In fact, the UNTB cannot be easily transferred in nature where a number of variables, such as environmental heterogeneity, biotic communities and coevolution move the debate in favor of the differentiation of niches. Anyway, in this paper I will show that neutrality can be considered as the basis for the subsequent separation of ecological niches.

## How much important predators and parasites are in the evolution of biodiversity?

The main issue that arises in relation to predation or parasitism and the evolution of the diversity of life is the ability of these functions to control the abundance of biological populations, change the reproductive rates and ensure a balance between the various species, moving it far away from an excessive dominance of a few that are common at the expenses of many that are rare. As these relationships are made of a network of different species and meta-populations, it seems almost impossible to define them with current mathematical methods. VanderMeer et al. (^26^) proposed a simplified model showing how the linear equations used in previous formulations, such as those of Lotka-Volterra, cannot take into account the real network of interactions that take place in nature between prey-predator/host-parasite. Their model allows, even if it is not able to describe all the dynamics, to understand how the competition among individuals is ultimately manifested in the coexistence of species. What seems clear, in spite of the many uncertainties, is that we cannot define - as we have done so far - the relationship of competition taking place for the direct purpose of alimentation, i.e. those involving the use of a prey by a predator or a parasite by its host to “gain energy”. This implies that we cannot consider predation, parasitism or grazing real cases of interspecific competition, as there is no membership at the same trophic level of the categories considered. We could instead re-evaluate the role of these functions as shapers of specific composition of ecosystems through the effects of top-down and bottom-up of predation and grazing, the variability in the diet or in the host, the presence of predators/parasites generalists and specialists, the mutual interference (^27^) and density-dependent effects (^28^,^29^).

Rather than competitive interactions for these categories (it would be better to define them as coexistence mediated by predators and mutualistic networks), this definition points out that all the interactions within food webs can allow the coexistence of many species in the same environment, due to their variability and complexity that influence the abundance of populations, the intraspecific competition and, consequently, the interspecific relations (^30^,^31^).

## The new hypothesis: how do we explain the current levels of biodiversity and species coexistence?

Recently, Levine and HilleRisLambers (^32^) have argued that niches are critical for the maintenance of species diversity, challenging the neutral theory of biodiversity which on the contrary explains the coexistence with the equivalence of competitors. This study, like many others, does not take into account the effects of history on biological diversity. A thorough understanding of the evolutionary dynamics of biodiversity, which could somehow explain the current distribution patterns and mechanisms of coexistence must consider the biogeographic and phylogenetic approaches. Only the union of geological, evolutionary and ecological dynamics in the context of natural history can show us a more clear and complete picture (^33^).

This can be done with a simple evolutionary graphical model of biodiversity (Fig. 1) that I made in the attempt to explain, using the latest scientific evidence, the mechanisms involved in species coexistence. I voluntarily avoided formalizing it in a mathematical model as our current knowledge and analytic techniques are still too simplified and embryonic compared to the natural networks.

**Fig. 1.**
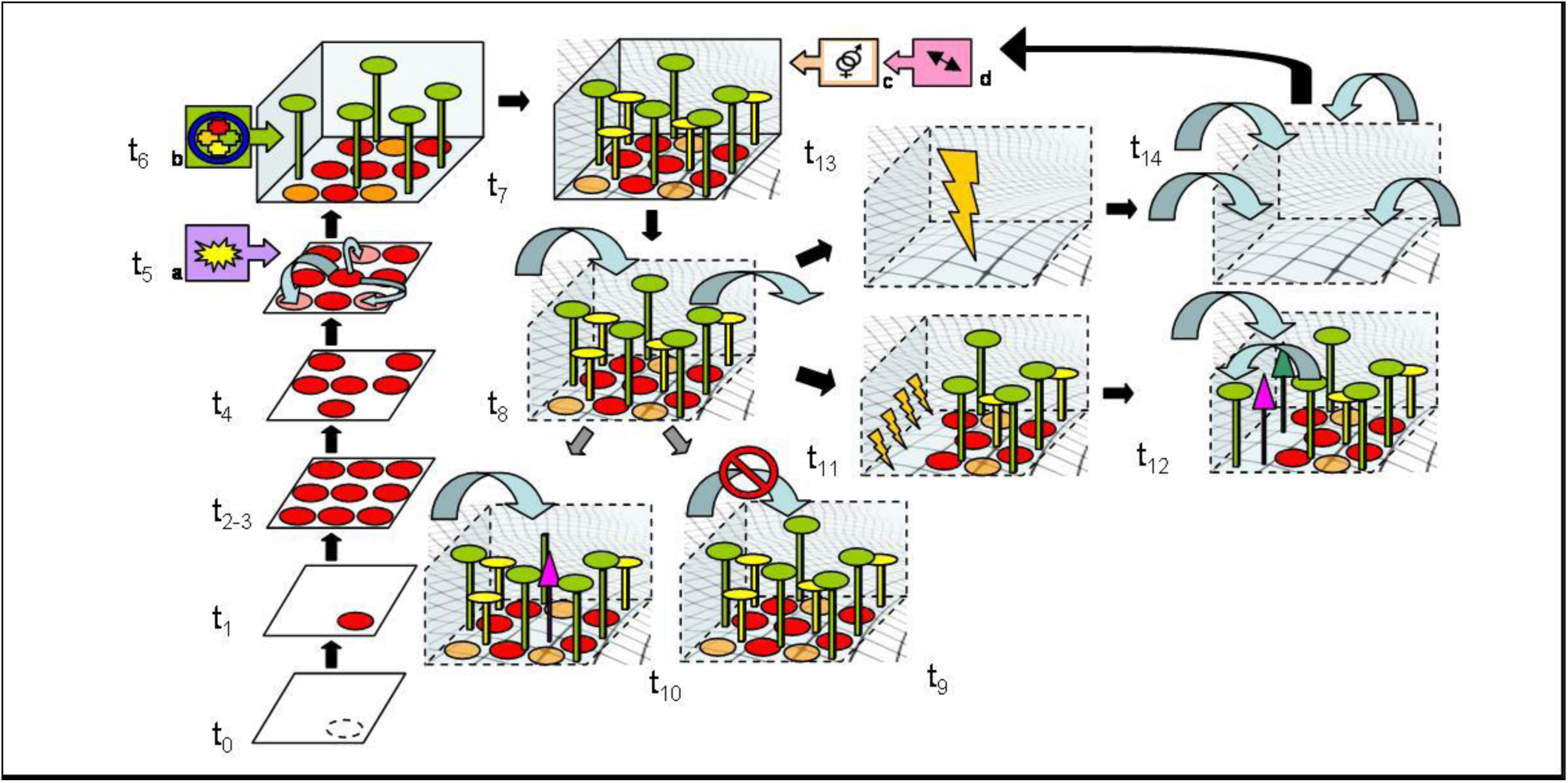
The evolution of biodiversity. Different steps towards species coexistence (see text for more details). t(0-14) are discrete times of the evolutionary process. Dotted-line circle is a potential species at t0. Red circles are a “two-dimensional” species. Pink circles are polymorphic meta-populations. Orange circles are new species. “Tree-shaped” species are those that exploited the 3-D niche’s environment. Grid landscape at t7 represents the heterogeneity of the environment. Dotted cube and blue arrows at t8 represents, respectively, the removal of barriers and the immigration and emigration processes. Small lightning at t11 are local extinctions and big lightning at t13 is mass extinction. a,b,c,d squares with arrows are respectively: virus and bacteria’s new alleles integration (*endogenosymbiosis*); BNDT effects including symbiosis, endo-symbiosis and mutualism; sexual reproduction and predation/grazing/parasitism events.

As Von Bertalanffy suggested in General system theory (^34^) “the advantages offered by mathematical models – non ambiguity, rigorous deductions, verifiability through data of observation – are well known. But this does not mean that models expressed with a common language should be rejected or despised. A verbal model is still better of no models or of a model that, being mathematically formulated, is forced to reality, falsifying it. […] The history of science attests that, frequently, the expression in common languages precedes the mathematical formulation, i.e. the invention of an algorithm”.

Let’s begin considering a two-dimensional environment (Fig. 1) where any form of life is lacking at time 0, which may be the Earth a few moments before the formation of the first living beings. Imagine, for the moment, that this area is isolated by a physical barrier (island, mountain chains, acidic waters, closed pool, etc..). The first cell that will be formed at time 1 will probably consist of a single individual that through asexual reproduction, when resources are available, will reproduce itself exponentially until time 2. While the amount of time before the formation of the first living organism is infinitely long (billion of years), the one between the first individual and the exponential increase of the population is relatively short. We can assume that all individuals that have reproduced so far belong to the same biological species. When reaching the maximum abundance allowed by the limited resources and carrying capacity of the environment at time 3, density-dependent effects (which will have an impact on birth rates and mortality) will take back the population within the limits of carrying capacity (time 4). In this simple environment, the death of individuals regenerates resources for the birth and development of other conspecifics.

After a relatively long time (million of years), it is possible that some integration of genome parts coming from successively evolved cell non-living parasites (virus and bacteriophages). These latter are just a combination of genes moving among cells and using them to replicate. Then the genetic sequences of the first species transcribe, leading to the formation of some individuals that are slightly or hugely (punctuated equilibrium^35^) different from their ancestors. We call them polymorphic meta-populations (^36^). When these differences are not so evident, to separate them reproductively (not only genetically but also behaviourally), we cannot define two meta-populations as two species. The population is constantly in a dynamic equilibrium that oscillates around the levels of carrying capacity. After many cycles of oscillations close to the limits imposed by the environment, one species cannot manifest a density-dependent effect reduction of populations. So it is possible that during a phase of exponential growth the species with the accumulation of new genes would fit - due to its phenotypic plasticity - to use a different resource from those used by the individuals of the original population (e.g. scrap products of it) or can use some resources more efficiently. Therefore in the first case it is the effect of a number of genomic inclusions, providing the characters displacement of individuals of the meta-population through phenotypic plasticity, that allows adaptation to different conditions. This process, defined as sympatric speciation, may occur by means of a shift in the fundamental niche of a meta-population from the original one (time 5). Recently it has been argued that true sympatry may not exist in nature. This is because small variations in the microhabitat preference can still create allopatry and recent investigations in habitat suitability studies seem to reveal these differences (^37^, Rissler & Apodaca 2007, Systematic Biol 56(6): 924-942^38^). If we consider sympatry as a spatial variable, the “microhabitat preferences” are not properly sympatric but instead represent a niche displacement. I suggest that sympatric speciation should be reconsidered as one of the main mechanisms that lead to species coexistence and to the evolution of biodiversity. In fact, if interspecific competition and the principle of competitive exclusion between meta-populations and the different lineages were to take place, probably there would never be the formation and coexistence of different species, but rather the survival of the most efficient one (that accumulates enough mutation to adapt) and the extinction of the ancestor or those belonging to other phyletic lines. The coexistence of two species in a sympatric way can happen only if there is low competition or competitive exclusion between them, but rather a kind of *avoidance of competition* that leads to a slight shift of the niche of a meta-population that accumulated a series of genomic inclusions coming from other sources of genes (particularly viruses). Thus, eventually, it’s the avoidance of competition and the process that I will call *endo-geno-symbiosis* (hereafter the capacity of endogen “genes carriers” to share parts of their genome in a symbiotic relation) that shows the ability to produce the diversity of life. Competition and mutation, on the other side, lead to preserve and adapt species and not to diversify them. This confirms what has previously been pointed out: in reality you cannot attend the competition in the present since all niches of the relevant species in an ecosystem seemed to be unique and different though overlapping in varying degrees. This is possible since there has never been competition between two species of the same territory because of their common evolutionary history and phylogenetic.

Continuing with the evolution of the system (time 6), further species formation is achieved by characters displacement and the realized niches through the avoidance of competition and endosymbiosis^39^. This can lead to reach again the limits of the carrying capacity with minimum viable populations of many species. The ability of species to change their external environment (imagine the production of oxygen), to develop in a three-dimensional space (and not only in two dimensions) and create relationships which take advantage of symbiosis and mutualism, may allow the expansion of the basal hypervolume and the formation of new species (even multi-cellular). At this stage the facilitation process plays a major role, that is to say the process that allows the development of new species taking advantage of the presence of others (hypervolume expansion) (^40^).

Going further (time 7) we see how the interaction between the biotic and the abiotic component increases the spatial heterogeneity of the ecosystem and this encourages further development of species that adapt to the new possibilities (formation of chemical elements, erosion of rocks, biochemical changes, etc.). Obviously, at the same time, it is likely for some species not being able to survive to the changing of external conditions to become extinct, consequently making niches available for speciation of others. The possibility of extinction due to environmental effects is, thus, more concrete than the one due to interspecific competition (^31^). Certainly at this stage, interactions of predation, grazing and parasitism between species will be originated. It will maintain, through the principles discussed above, the various populations in dynamic equilibrium. This situation could persist for a long time or change after the removal of environmental barriers hitherto considered (shift of continents, climate change, drying up of rivers, etc.). At the time 8, the species whose populations fluctuate around the carrying capacity will, by chance or movements to reduce the effects of density and, therefore, aimed to avoid competition, migrate to neighbour territories and, thus, immigrate into new areas. Here we can find two possibilities. The first is that the environment into which these individuals migrated does not have available niches so, as a consequence, they will be rejected (time 9). The second is that the immigrant species finds a niche available and settles in that territory (time 10). More and more evidence (^41^) shows that in intact ecosystems the chance that an alien species can survive is very low and that ecosystems under stress are more prone to having available immigrant niches to let alien species settle in. Emigration can either still keep the meta-populations in contact and thus genetically belonging to the same species or, because of the distance and/or the rebuild of physical and geographical barriers, it lead to the formation of two distinct species (parapatric/allopatric speciation). Allopatric speciation then, although it can occur suddenly for the formation of physical barriers, is less likely to appear than sympatric as the result of inclusion of new parts of genomes and phenotypic adaptation to new niches. Because either in the short or long-term environmental conditions tend to vary, it may happen that ecosystems that face low and time-limited instability see some species become extinct because they do not adapt to those conditions (time 11). These areas have a high rate of immigration since some niches remain empty due to extinction. Even in this case, it will be very unlikely for competitive exclusion between immigrant and local species to occur, since immigration is dependent on the distance (^16^). It is more likely for a species of neighbouring territories, and therefore phylogenetically close to those extinct, to emigrate filling the niches gap without potential competition (time 12).

In case of environmental instability of greater intensity or duration the extinction of species in a given ecosystem is likely to be massive and that, therefore, the rate of the immigration of species from the surrounding environment, in an attempt to reoccupy the many empty niches, is very high (time 13). Certainly the need of speeding up the ability to face variable environmental conditions and the appearance of parasites has encouraged and allowed the evolution of sexual reproduction. This latter through the recombination of two different genetic pools and the increase of the probability of favourable mutations adapt, but not change (even in the long terms), the species to new external conditions (^42^,^43^). Such a change results from a compromise between the need to transmit in a long-term as much as possible genetic material to future generations and the need to address the environmental variability, while halving its fitness potential. In this perspective, the evolution of sexual reproduction appears as an extreme action to preserve the species, in other words to adapt them to external changes and not, as hereto always suggested, as a mechanism capable to produce new species due to mutations. It is furthermore likely that the same cause of the evolution of sexual reproduction, the parasitism, represents also the beginning of biodiversity. In other terms, it seems that sexual reproduction acts as a conservative system against the inclusion of new genetic variations into cells’ DNA (supported by the mutations reparation systems) and, instead, the evolution of species appears only when this preservative system fails to contrast the inclusion, within the host genome, of hexogen parts of DNA coming from “parasitic” cells (viruses, phagi). As two parallel evolutionary lines, sexual reproduction seems to preserve what the *endogenosymbiosis* moves to diversify. Following the first, the species can adapt slowly and indefinitely to the external factors, adjusting themselves (adapting) but not “creating” novelty. The second line leads to speciation due to sudden changes in genes sequences.

The dynamic process just described may, therefore, continue to be used for further endless cycles of sexual reproduction, while maintaining the basic patterns set.

These dynamics, considered in biogeographycal and evolutionary terms, seem to be the most likely to have led to biodiversity and the coexistence of the species that we see today on our planet, both in aquatic and terrestrial ecosystems.

## The evolution of biodiversity in microscale: the importance of the *avoidance of competition* and cooperative mechanisms

The simplified model of the evolution of biodiversity described so far leaves out two aspects of microscale that cannot be omitted in a comprehensive view of the processes that affect species diversity.

In particular, it is necessary to investigate what happens at time 3 at the maximum carrying capacity of the only species present or at time 5 with different species (Fig. 2). Exceeding the carrying capacity by some populations, in case of no possibility of emigration even on the microscale, leads to the appearance of a series of density-dependent effects (mutual interference, resource depletion, diseases, grazing, predation), which bring them under the limits allowed by the ecosystem (increasing mortality rates rather than the birth). According to the statement of the principle of competitive exclusion, when a balance between population/resources is reached and all individuals of the same species obviously have identical niches, there can be no way for the expansion of the species or for development of new ones. What actually happens, however, follows precise steps that lead to the increase in the number of species.

**Fig. 2.**
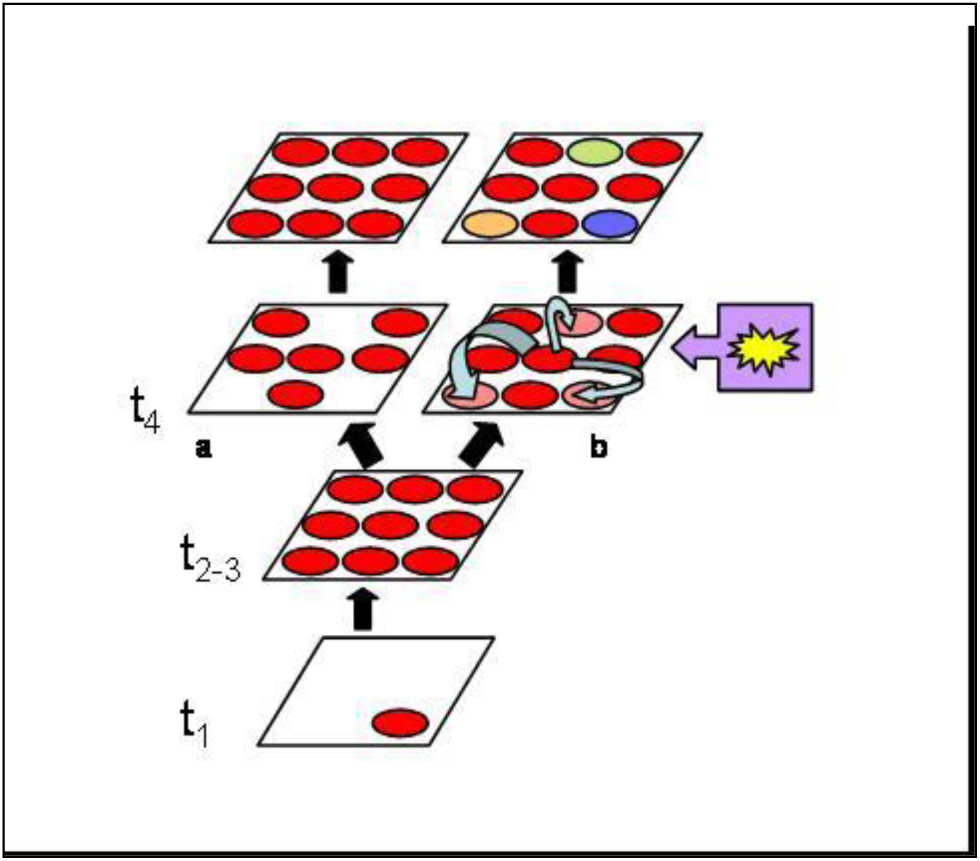
The avoidance of competition leads to species coexistence. At t4 the empty space due to cyclic variation of population around the carrying capacity is filled by individuals of the same species (a), with mutation that allow them to adapt but not to evolve in different species. (b) with *endogenosymbiosis* and phenotypic plasticity these latter (purple square with arrow) some polymorphic meta-populations can evolve in different species following the principle of the “avoidance of competition” (see text for more details).

A series of random mutations induced by either *endogenosymbiosis*, environmental or reproductive (less frequent if asexual than sexual) causes may lead to an accumulation and a change in genotype of some individuals of the species (characters displacement and polymorphism). This will generate a phenotypic plasticity that can be used to shift slightly the niche and avoid competition for space and resources with non-mutated (wild type) individuals. In this way new species can be formed (following the accumulation of behavioural differences as a result of niches shifting and then sympatric reproductive isolation), which coexist in the same territory as they have different needs. It is well known, for example, the case of the coexistence of wild dogs, lions and jackals whose ancestors of different phylogenetic lines, during their evolutionary history, established trade-offs in the use of the same prey, but in different ways, ensuring the survival in the same territory, precisely through the avoidance of competition.

Something similar happens in the plant world, where we can find (especially in tropical forests) many species with apparently the same ecological requirements, whose niche were slightly differentiated from their phenotypic plasticity, driven by the avoidance of competition, thus ensuring the coexistence of species that are apparently very similar.

The avoidance of competition, therefore, not only resizes the importance of competition in relationships among species, but also explains why steadier ecosystems with intermediate disturbances show greater levels of biodiversity (as in the case of coral reefs and tropical forests). These steady ecosystems are able to iterate the cycle of the model described above, minimizing extinction and increasing speciations. Temperate forests, which have experienced recent phases of glaciations with high climatic variability and deep stress because of anthropogenic activities, have reset many times the cycles of “speciation through the avoidance of competition” leading to numerous extinctions and, therefore, they now include far fewer species than tropical.

A second mechanism of microscale that we must consider to complete the framework of discussion is what happens at time 6, when the species begin to exploit the three-dimensional space (Fig. 3). Both in laboratory experiments and those in the field, the three-dimensional space of biodiversity is often deliberately omitted, conducting studies that force species to interact in a two-dimensional environment. In fact, the three-dimensional space provides a greater exploitation of the niches made available by the environment.

**Fig. 3.**
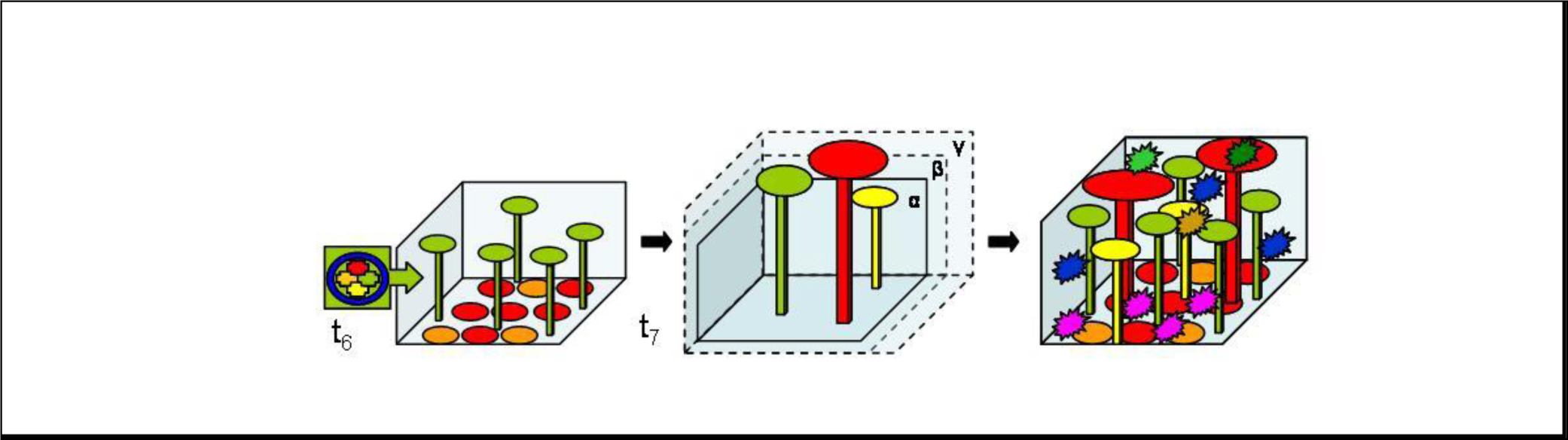
Species exploit the 3-D space and the BNDT effects. As suggested by the BNDT the exploitation of 3-D space and the species facilitation allow the increase of basal niche space and enhance the number of species that can fill the ecosystem (see text for more details). Green square with arrow represents the mechanisms foreseen by the BNDT. α, β, γ cubes are different basal niches shaped according the environmental availability (in particular temperature and humidity). α could be a polar niche with limited space and few species, β could be a temperate ecosystem and γ a tropical ecosystem with higher temperature and humidity and, consequently, bigger 3-D niches space that key species can exploit allowing more (as in the last cube of the figure) animal and plant specialist species to survive.

With the Biodiversity-related Niches Differentiation Theory BNDT (^35^) I recently proposed that species themselves are the architects of the greatest biodiversity of a given environment because, through the realization of their fundamental niche, they allow the expansion of the available niches for other species. This is not a form of the species-area curve (e.g. more tree surface relative to shrub is equal to more species) but a new theory to explain species coexistence. Although every species is able to support the establishment of many others, it is especially the generalist and structuring ones (key-species) that greatly increase the hypervolume available in the ecosystems. They do this by extending the three-dimensional space that allows the specialist species to establish themselves in the same habitat (in height or in depth). According to this theory, phenomena such as overvalued intra and inter-specific competition, gives way, reconsidering their real importance in the evolution of biological diversity, to mechanisms such as facilitation, mutualism, symbiosis, coevolution, which all lead to the coexistence of species.

The BNDT also acts as a glue between the theory of neutrality and the one of the specificity of species and of the niches they create. According to the BNDT, generalist species expand the basal two-dimensional hypervolume (with a limited number of niches available) through the creation of three-dimensional niches. Their lack of specificity and their high equitability follows the provisions of the neutral theory. Once created by generalist species, the niches are filled (speciation/immigration) by specialist species that show differentiation, following the principle of the avoidance of competition and stability, adhering to the requirements of the theory of niches differences. Therefore, the evolutionary dynamics of ecosystems follow the development of biodiversity ranging between the equitability/neutrality of early successional and the specificity/stability of lately stages. Between them the effects of the BNDT occur.

A final, but important question that can be solved by BNDT concerns the problem of why tropical ecosystems are the richest in biodiversity and why ecosystems that receive more energy have got more animal species. Based on the predictions of BNDT, tropical ecosystems receiving a greater amount of light energy (thus temperature) and rainfall (thus humidity) have a broader basal hypervolume than temperate ones. Without taking the effects of BNDT into account, we might mistakenly conclude that the high amount of productivity and available resources in regions with much more energy does not explain high biodiversity in these areas as there could be only a high rate of available resources and not a greater variety of these, such as to justify the higher number of niches and species. The highest three-dimensionality (volume of niche basis) due to higher energy is even at the base of the explanation of why some ecosystems contain more animal species. If you imagine, for instance, an ecosystem like this or polar tundra with a sparse and mostly distributed in a two-dimension vegetation, compared to an abundant tropical vegetation and especially developed in three-dimensions (think about the enormous trees with buttresses, the vines, the epiphytes, etc.) it is easy to understand how both the amount of energy and the facilitation due to the implementation of biodiversity-related niches, allow the basal hypervolume to expand and accommodate many more animals and plant species. It is easily proved that a creeping shrub of Siberia will host a number of limited animal and plant species compared to a tree in Borneo. The temperate ecosystems are placed midway between these two extremes. The effect of higher dimensionality of hypervolume could be, finally, an explanation of why temperate ecosystems contain fewer large mammals than tropical ones.

## Conclusions

Much emphasis has been put on the competitive interactions between individuals of the same species and different species in ecological and evolutionary studies rather than on cooperative approaches. Darwin himself wondered how the simultaneous presence of many species in one ecosystem could be justified in a struggle for existence. The allopatric speciation has often been cited as the main cause of species formation, underestimating the importance of the mechanisms of sympatric divisions. Recent researches and developments of new theories on the mechanisms of cooperation and facilitation between individuals (^44^) and species (BNDT) and the role of sympatric speciation (^45^,^46^) must be significantly re-evaluated as the main factors in the evolution of biodiversity.

Critics of this approach may highlight the fact that many alien species that have been introduced deliberately or accidentally into territories far from their habitat in recent decades, contradicting what hereto argued, have developed competitive behaviour up to eliminating or greatly reducing native species with their similar realized niche. However further analysis is needed to observe two elements. The first is that alien species are rarely able to settle in areas not subjected to anthropogenic stresses (alteration, fragmentation, selective logging or hunting, etc.) and that when an allochthonous invasion takes place, it is very unlikely for the alien species to completely eliminate the endemic one, which usually tends to reduce their density or migrate to neighbouring areas, when it has the phenotypic plasticity to do it. In rare cases, resource and habitat limitations may lead to the extinction of native species (^47^). This latter event, however, cannot be considered as an effect of the principle of competitive exclusion. The two species that come into contact in this way, usually carried by human activities over long distances, come often from different continents whose biological histories and phylogenetic relationships are so far away and generated through allopatric parallel evolution with identical niches. This suggests that the evolution of biodiversity itself has the tendency to avoid competition (the principle of *the avoidance of competition*). Competition can never happen between two species that come from a common biological and phylogenetic history and can occur only in case of deep alterations of the ecological balances (stresses and invasion of alien species). In invasion events the two species in contact, being distant branches of the evolutionary tree, have not been able to establish a series of trade-offs which might prevent competition and have to interact when their coexistence is forced.

In nature, therefore, competition seems to be the exception and the avoidance of it by any means the rule (even in intraspecific relationships we talk about rituals put in place to prevent damages caused by competition for resources, space or mating). To prove this latter statement, in addition to the the evidence already reported, there is a large number of defence mechanisms (poisons, spines, allopathic substances, dispersal in plants, animals migrations, etc..) evolved in order to avoid competition between species belonging to the same or different trophic guilds and among individuals of the same population. Those mechanisms who have been interpreted so far as competitive are in reality nothing more than the systems used to avoid competition, save energy and, consequently, allow the coexistence of species with similar niches. Thus the evolutionarily stable equilibrium (ESS) in any case is not the competitive, but the avoiding-facilitative one.

These conclusions have often been criticized for their lack of mathematical evidence able to model these laws. But we must admit that the complexity, non-linearity, feedback mechanisms make the networks of relationships in an ecosystem virtually impossible to be represented with the current mathematical tools. The same stability against stress of more complex systems has been questioned (^48^) by mathematical models, against the empirical evidences. This may be due not to the fact that ecosystems with high biodiversity are really more vulnerable but to the evidence that non-linear mathematical models with many variables are not easily manageable with current analytical techniques and therefore tend to generate erroneous results.

Finally, criticisms to the old hypotheses and ideas suggested here by the new developments recall the urgent need to halt biodiversity loss caused by human beings. Marine and terrestrial ecosystems are subjected to efforts that do not match with their ability to regenerate. The great extinctions of the past, such as the Permian-Triassic and the Cretaceous-Tertiary, were all caused by climatic and physical-chemical changes (^49^,^50^) and not by competition for resources and space between species. This should push our species to think before it is too late about how human competition, for the first time in the history of life on Earth, has been leading to the extinction of animals and plants systematically. The simple model of evolution that I proposed here does not only explain the mechanisms that underlie the current presence of myriad forms of life but it also sheds new light on the need of periods of geological scale time (evolutionary mechanisms) rather than periods of biological (ecological mechanisms) to generate the awesome number of species that currently inhabit our planet. If humanity does not stop its “unnatural” competitive spirit in the massive elimination of species, it will take thousands of years before the diverse set of life forms, that we now call biodiversity, will be regenerated.

